# Speech in noise performance in adults with cochlear implants using a combined channel deactivation strategy with a variable number of dynamic focused channels

**DOI:** 10.1101/2024.10.31.621419

**Authors:** Dietmar M. Wohlbauer, Charles Hem, Caylin McCallick, Julie G. Arenberg

## Abstract

**Objectives and Methods:** Cochlear implant listeners show difficulties in understanding speech in noise. Channel interactions from activating overlapping neural populations reduce the signal accuracy necessary to interpret complex signals. Optimizing programming strategies based on focused detection thresholds to reduce channel interactions has led to improved performance. In the current study, two previously suggested methods, channel deactivation and focused dynamic tripolar stimulation, were combined to create three cochlear implant programs. Utilizing an automatic channel selection algorithm from focused detection threshold profiles, three programs were created with the same deactivated channels but varying proportions of channels employing focused stimulation, monopolar, dynamic focused and a mixed program. Thirteen ears in eleven adult cochlear implant listeners with Advanced Bionics HiRes90k devices were tested. Vowel identification and sentence perception in quiet and noise served as outcome measures, and the influences of listening experience, age, clinical consonant-nucleus-consonant performance, and perceptual thresholds on speech performance were assessed.

**Results:** Across subjects, different degrees of focusing showed individual performance improvements for vowels and sentences over the monopolar program. However, only slight trends and no significant group improvements were observed. Focused listening benefits were shown for individuals with less cochlear implant experience, and clinically poor performers seem to benefit more from focusing than good performers.

**Conclusion:** The current findings suggest that deactivating and focusing subsets of channels improves speech performance for some individuals, especially poor performers, a possible effect of reduced channel interactions. The findings also show that individual performance is largely variable, possibly due to listening experience, age or the underlying detection threshold.

## 1 Introduction

Cochlear implants (CIs) are the most successful neural prostheses currently available to treat severe to profound hearing loss [1]. Nevertheless, CI listeners show difficulties in understanding speech in noise likely due to large channel interactions at the electrode-neuron interface (ENI). Channel interactions are the result of different channels activating overlapping neuronal populations [2]. Because each channel carries a different range of spectral information, channel interaction compromises spectral resolution, and CI listeners have difficulty with complex signals such as understanding speech in background noise [3–7].

Two methods to diminish unwanted channel interactions to improve perceptual outcomes in electric CI stimulation have been suggested previously. One method is to deactivate channels suspected of having a lot of channel interaction or a poor ENI. While several studies found performance improvements with a subsest of deactivated channels [2, 8–11], other studies did not [12, 13]. [11] investigated a clinical method to deactivate CI channels with channel interaction identified with indistinguishable pairs of electrodes in a pitch ranking task, which led to increased performance for consonant-nucleus-consonant (CNC) and AzBio sentences in quiet and babble noise Peterson and Lehiste [14], Spahr et al. [15]. In contrast, Vickers et al. [12] identified channels for deactivation in a similar experimental setup of pitch ranking but could not find improvements for sentences in noise. One possible underlying factor for the variable study outcomes was the different underlying programming strategy. While perceptual outcome benefits were observed with channel deactivation with the stimulation strategy that employs current steering, the main strategy for Advanced Bionics (AB) devices [11], both studies failed to show statistically significant benefits for channel deactivation with the Advanced Combination Encoder (ACE) strategy, the standard used in Cochlear^®^ devices. Hence, channel deactivation in N-of-M type channel selection might have less impact due to the variable selection of actively stimulating channels. A recent study tried to counteract the *N* -of-*M* channel selection by activating every other channel per stimulation frame, hence, by deactivating intermediate CI channels, and showed slight improvements in unilateral and bilateral speech perception [16]. These mixed results highlight the importance of selecting the correct candidate channels for deactivation, as well as the choosing the right speech processing strategy.

The second method to reduce channel interaction is current focusing, which has shown a reduced spread of neural activation at the ENI in animal models compared to the standard monopolar (MP) configuration used clinically [17–21]. Focused stimulation incorporates a biphasic cathodic leading stimulus on one electrode accompanied by a simultaneous biphasic anodic leading stimulus on return electrode(s) within the cochlea. The tripolar configuration returns current to the neighboring electrodes to both sides of an active electrode. This method could be more similar to the narrow activation along the basilar membrane by pure tones in normal hearing (NH) [18, 22]. However, there are several limiting factors to using tripolar in speech processing strategies, such as the difficulty to generate a sufficiently loud signal per tripolar stimulation complex, and a relatively high power consumption [23, 24]. Previous studies show that current focusing requires higher stimulation currents to generate a similar loudness percept to monopolar stimulation [23–25]. In addition, the higher stimulation currents increase the implant’s power consumption. To counteract these drawbacks, partially tripolar (pTP) stimulation was implemented, such that instead of returning all of the stimulation current via the flanking electrodes, a fraction of the current returns via an extra cochlear ground electrode[2, 17, 25–31]. The studies that implemented the pTP configuration in speech processing strategies observed some improvements on spectral resolution tasks and speech perception [2, 29], but there were also some mixed results [26].

To approach the variable outcomes of studies using tripolar and pTP, the method of focused stimulation was extended by a physiologically inspired idea based on a loudness model by [32]. In the so-called dynamically focused or dynamic tripolar (DT) stimulation, the proportion of focused current of the pTP stimulation is dynamically adapted based on the stimulation level [33, 34]. This procedure mimics cochlear excitation patterns, where low-intensity signals are more focused on a neural distinct area than high-intensity sounds, that excite a broader area. Consequently, a sufficient signal loudness can be provided, and the current needed to generate a loud signal is reduced; hence, power consumption could also be improved. The DT strategy showed consistent improvements for vowel identification in noise over the MP and the pTP strategies [29, 33, 35], and for individuals for whom ENI likely contributed to the benefit observed with DT.

Previously, one study investigated a combination of both methods, deactivating and focusing electrodes in one strategy [2]. Electrodes for deactivation were selected similarly to previous work via a poor ENI, a relationship identified to reduced performance due to higher perceptual thresholds and increased channel interactions [6, 36]. To further improve channel selectivity, the goal of the study was to create the highest possible degree of focusing on the remaining channels, respectively, the channels with a good ENI, defined via low perceptual thresholds. The study showed, that clinically ”poor” performers improved with pTP stimulation in a reduced set of active channels compared to MP stimulation. Nevertheless, perception scores for focused stimulation with a subset of deactivated electrodes did not improve for the whole study population and were highly variable.

Optimizing programming for high-focused threshold channels in DT and low channels in MP might improve speech perception in noise, similar to the previous findings with pTP stimulation. Addressing large threshold jumps from one channel to the next by manually deactivating or focusing might, therefore, be crucial to optimize individual stimulation. In addition, the DT stimulation could reduce power consumption compared to traditional pTP and power could be further reduced by stimulating only a set of high threshold channels in DT, and a set of low threshold channels in MP (i.e. some MP instead of DT channels, and less active channels result in less required power).

An overarching conclusion from the previous studies, is that there is still room to develop individual program optimization approaches, with a next step of combining channel deactivation and dynamic focusing some or all remaining channels. Hence, the present study investigates the performance of individually optimized dynamically focused programs to improve speech perception outcomes in quiet and background noise. The current study creates experimental programs that have a subset of channels deactivated based on high focused threshold profiles, either with the remaining channels in monopolar (MP program), a subset of relatively high threshold channels in DT (Mixed program), or all remaining channels dynamically focused in DT (DF program). Speech perception scores were obtained in quiet and noisy background listening situations for medial vowel identification and sentences. The current work aims to assess if the Mixed and DF programs perform better in sentence recognition and vowel identification tasks in quiet and background noise than the experimental MP control program. In addition, outcomes will be analyzed in terms of variables derived from the subjects’ demographics and individual detection threshold measures.

## 2 Methods

### 2.1 Subjects

Thirteen ears in eleven postlingually deafened adult CI recipients (two bilaterally implanted) between 44 and 76 years of age (average of 62.7 years), who were using their CIs between 1 and 24 years (average of 9.6 years), participated in the presented study. All participants were native English speakers implanted with the Advanced Bionics HiRes90K device and used HiRes Optima-S as their everyday coding strategy. Clinical pulse rates/width ranged from maximal 1856 *pps*/18 µs to 932 *pps*/35.9 µs. AB’s ClearVoice (Buechner, 2010) was set to “Moderate” in all participants except for A002, who had ClearVoice set to “Off”, to match their clinical programs for both ears. The clinical input dynamic range (IDR) setting showed similar settings throughout all participants (70 dB ± 5 dB). Participants 65 years and older completed the MoCA test and those scores, demographics, etiologies, and the clinical program parameters are shown in Table 1.

**Table 1:**
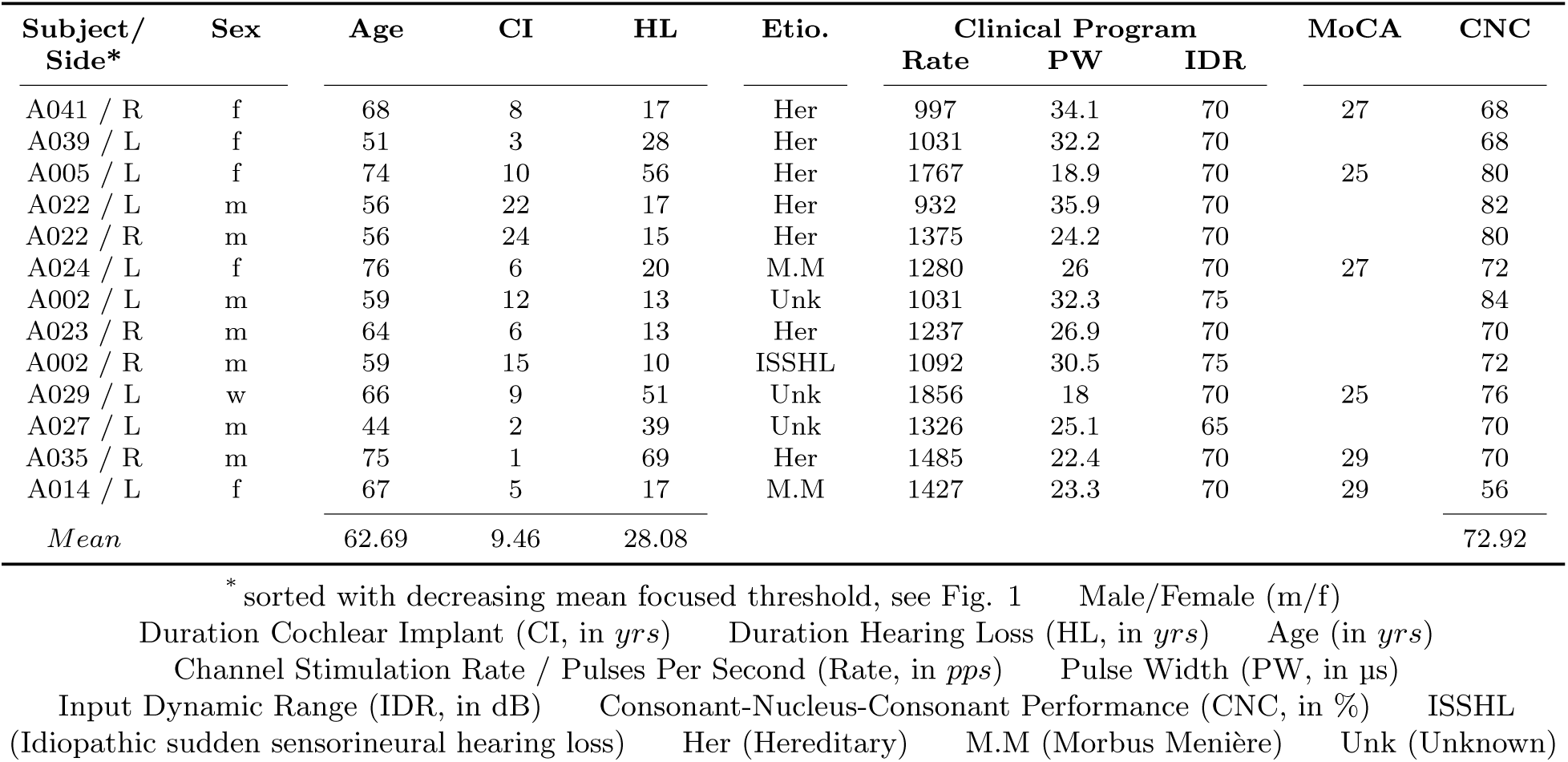
Study participant demographics and clinical program.

### 2.2 Focused threshold levels

Focused threshold recordings were performed in all thirteen ears to identify their detection thresholds and were used for determining channels to deactivate and/or focus for the experimental programming and the analysis. The procedure consists of several steps; To set up the upper stimulation limits for the focused threshold measurement, the most comfortable loudness (MCL) was obtained behaviorally for active electrodes 2 to 15. Subjects were asked to rate loudness using AB’s clinical loudness rating scale as the level was slowly increased, and to indicate the current level until loud-ness 7, “loud but comfortable”. Then the level was reduced to the loudness rating of 6 or “most comfortable”. When MCL was found, focused thresholds were recorded with a fast psychophysical sweep procedure (Bierer, 2015). Custom-made MATLAB software was used to control the experiment and stimuli via AB’s Bionic Ear Data Collection System software (BEDCS version 3). The sweep procedure uses a steered quadrupolar (sQP) electrode configuration. In sQP a fraction of the return current, defined via sigma (*σ*), is delivered to the flanking electrodes, and the remainder is delivered to an extracochlear ground. A value of *σ* = 0.9 was selected in this study to maintain current levels lower than the voltage compliance limits of the device. A second steering coefficient alpha (*α*) controls the steering between the apical and basal electrodes, with *α* = 0 to deliver the stimulating current to the apical, and *α* = 1 to deliver the current to the basal electrode. Pulse trains are presented at regular intervals while the *α* value is increased from 0 to 1 in 0.1 steps from the most apical to the most basal set of electrodes. This process repeats continuously as a forward sweep along the array until all possible sets of electrodes ranging from electrodes 2 to 15 are assessed. The same sweep is repeated in a backward manner and the whole procedure is repeated twice before being averaged together which comprises a complete threshold measurement to estimate the focused perception threshold profile for each participant, shown in Fig.1.

**Figure 1:**
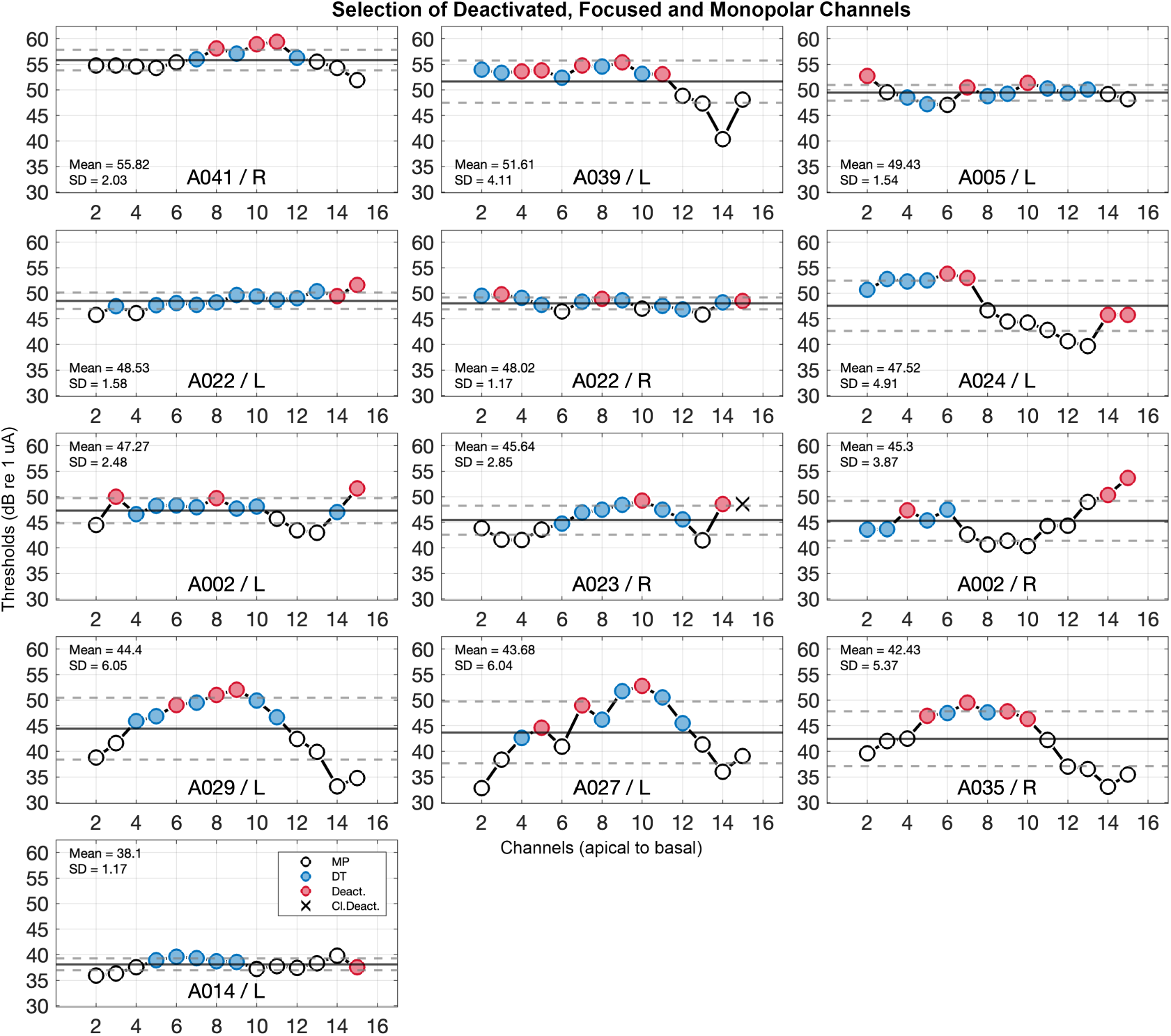
Focused threshold profiles measured with the fast sweep procedure (Bierer et al., 2015). Indicated are the optimized selection of deactivated and focused channels as red and blue circles, and the remaining MP channels as black circles. Participants are sorted from high to low mean focused thresholds; mean and ±1 dB re 1 µA SDs are indicated as solid and dashed lines. Subject (A023/R) had channel 15 clinically deactivated, indicated with (X).

### 2.3 Channel selection for experimental programs

Based on the focused threshold profiles, three experimental programs were created for this study using AB’s BEPS+ research platform and implemented on a Harmony research processor. The three programs were such that active electrodes were: 1) programmed in MP, 2) programed with a subset in MP and DT (Mixed), and 3) programmed with all remaining channels DT (DF). All three programs had the same subset of deactivated channels, including channels 1 and 16 given the restrictions of focused stimulation with one reserved flanking channel on each side of the stimulating electrode. The programming software optimizes the power consumption, due to higher power requirements for DT channels (i.e., automatic adjustment of rate and pulse width according to the required charge, lower rates allow longer pulse durations and lower amplitudes and result in equal charge with a lower power requirement). The participants’ experimental IDR and ClearVoice settings were matched to their clinical settings (Table 1) throughout all assessed experimental strategies. The detailed parameters of the experimental programs are listed in Table 2.

**Table 2:**
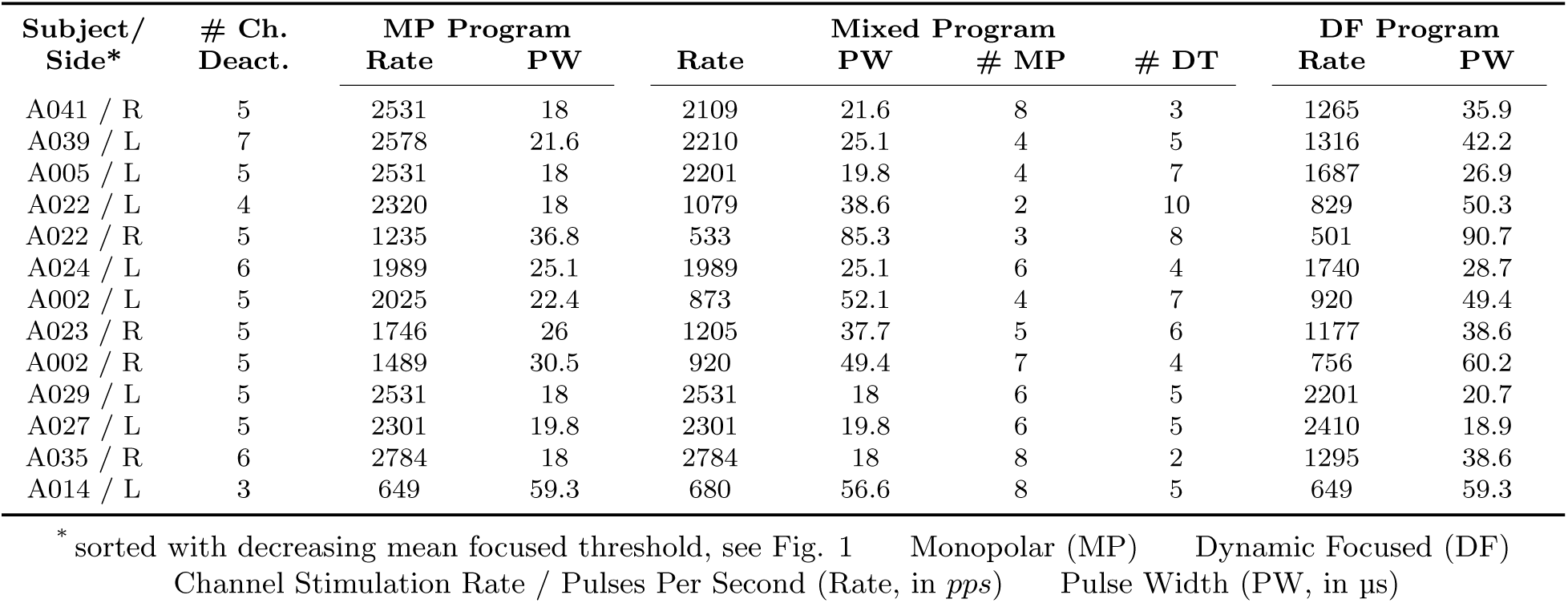
Experimental program parameters.

Bierer and Litvak [2] presented a method to automatically select channels for deactivation, which inspired the channel selection approach applied here. The initial method was to deactivate high-focused threshold channels to reduce the standard deviation (SD) of the enhanced profile until a subject-specific criterion or at least eight active channels remain. The profile was enhanced to increase the contrast of the threshold variability, so that high and low threshold channels could be clearly identified with the following restrictions: 1) maximally deactivate six channels, and 2) maximally deactivate two channels in a row.

To optimize the automatic channel deactivation for the three experimental programs of the current study, the suggested method by Bierer and Litvak [2] was revised. To pre-process the profile peaks and troughs, a contrast enhancement was performed, similarly to the previous study, however with an increased gain of six instead of four, to further improve the contrast and more clearly distinguish the individual profile variabilities. High threshold channels were deactivated based on the individual average threshold and the limit of +1 dB re 1 µA SDs, when channels of the enhanced threshold profile exceeded that limit. The algorithm was further limited to maximal two deactivated channels in a row, to avoid the deactivating too broad of neural regions. If more than three neighboring channels were selected for deactivation, the lowest channels were excluded from the subset of deactivated channels or changed to DT in the Mixed program. The subsets of deactivated channels are shown in Fig.1 as red circles.

With channels defined for deactivation, the next step was to select which channels would be focused and which would remain monopolar, shown in Fig.1 as blue and white circles. In addition to the Mixed channels automatically assigned due to the deactivation restrictions, see above, channels selected for focusing for the Mixed program were chosen if their enhanced threshold was between the threshold average and the applied +1 dB re 1 µA SDs criterion, while channels below the average threshold were set to MP. To finalize the experimental programs, the subsets channels were visually inspected, and the selection was individually revised for an optimal individual setup. In some cases, the automatic channel selection did not properly detect candidates for deactivation and DT stimulation, especially when the profiles show high variabilities or a non-symmetric tilt as the result of either high or low thresholds to one side of the profile. If neighboring threshold channels showed large variabilities, but the enhanced profile did not clearly highlight the channels for deactivation by exceeding the mean by +1 SD, or for setting them to DT, the candidates were manually selected. The final channel selection is indicated in Fig. 1, with red circles (deactivated channels), blue circles (DT channels), and black circles (MP channels). In addition, clinically deactivated electrodes are marked by (X).

Validation of the consistency in the channel selection procedure across subjects, represented as mean-corrected threshold levels in Fig. 2, shows, that the highest threshold channels are consistently selected for deactivation and the lowest for MP. The subset of channels set to DT stimulation for the Mixed program are generally above the individual threshold averages, while MP channels show values below the average. The selection of two study participants, resulted on average in higher focused than deactivated threshold channels, due to their high variability in focused thresholds of deactivated channels (A024/L), due to the mostly flat threshold profile and overall close threshold levels (A014/L), or the general limit of two deactivated channels in a row, which might lead to additional focused channels at high thresholds.

**Figure 2:**
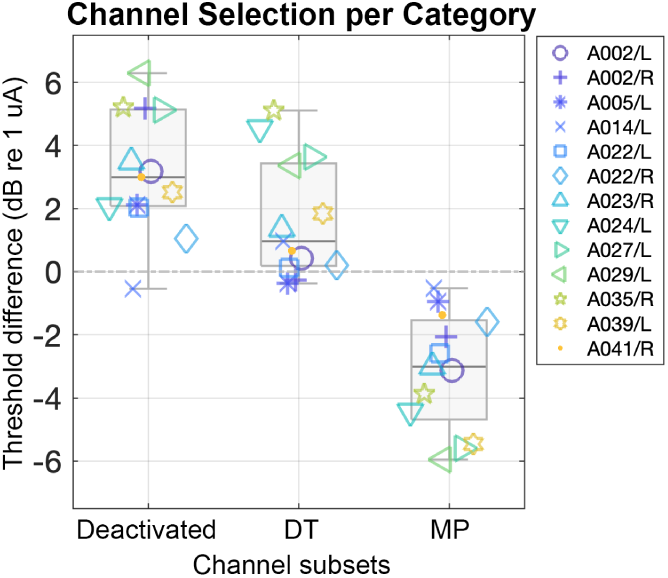
Mean-corrected averaged thresholds of CI channels selected for deactivation, and DT and MP channels of the Mixed program.

### 2.4 Programming

After defining channels for deactivation and focusing, the three experimental strategies were implemented in BEPS+. MCLs and threshold were measured with the same behavioral procedure that was used to find focused MCLs for all channels as described above. Pulse trains were presented starting at inaudible levels and were increased until the participant detected a sound. Then levels were reduced by 5 CUs. Stimulation levels were then increased to reach MCL. Following these settings, loudness balancing was performed at MCL with sets of 4 channels to ensure equal loudness. Finally, the strategies were switched to live mode for additional fine tuning as needed. An additional stage of acclimatization included 15 minutes of listening to AzBio sentences with each strategy [15]. Two participants (A029 and A035) experienced disturbing background noise in the DT programs. Lowering threshold levels reduced the noise percept to some extent, however, this was not optimal and not comparable to the other participants therefore those participants did not complete the testing with those strategies.

### 2.5 Outcome measures

Speech perception testing included sentence recognition with open-set IEEE sentences and closed-set vowel identification test with ten naturally spoken male-talker medial vowels in /hVd/ context (heed, hid, hayed, head, had, hod, who’d, hood, hoed, hud). Speech testing was performed in quiet and in the presence of Auditec four-talker babble background noise. Speech stimuli were presented at 60 dB SPL and the background noise level was adjusted to estimate signal-to-noise ratios (SNRs) for approximately an easy (80% correct words) and a difficult noise condition (60% correct words). Two lists of 20 sentences were presented at each SNR, and scores for all four lists were averaged together and are described as a noise condition. Sentence and vowel SNRs for the noise conditions were determined for each participant individually, using their clinical baseline strategy with automatic noise-reduction (AutoSense 3.0) off, to best match processing available in the Harmony research processor. Note that an SNR of +25 dB was the limit of our system, therefore this was used for participants’ easy condition (SNR80, even if they could not achieve 80% correct) and a slightly smaller SNR was then used for a more difficult condition, see 3 for the SNRs used for each participant.

Testing was performed in a sound-treated booth, and via a tablet PC (Windows Surface Pro 7, Windows 10) and custom software. Study participants were seated at a table with the tablet placed at a comfortable distance. The sentence lists were presented acoustically through the tablet’s speakers at 60 dB SPL in randomized order so that the individual lists were only presented once. A total of six lists were presented, two training lists with 20 sentences, and two test lists per SNR with 60 sentences. Participants were instructed to repeat the heard sentence, which was recorded. Recordings were scored for correct words, outcomes averaged, and converted to percent correct. The vowel lists were presented the same way as sentences but with the words shown on the tablet’s screen. Participants were instructed to choose the heard word from the available list. Each word was repeated six times, and the outcomes were averaged and transformed into percentage correct words.

**Table 3:** Study participant sentence and vowel SNR levels.

| Subject/<br>Side | Sentence Recognition |  | Vowel Identification |  |
| --- | --- | --- | --- | --- |
|  | SNR80 | SNR60 | SNR80 | SNR60 |
| A041 / R | 25 | 20 | 20 | 17 |
| A039 / L | 25 | 23 | 13 | 10 |
| A005 / L | 20 | 18 | 10 | 8 |
| A022 / L | 20 | 16 | 7 | 4 |
| A022 / R | 20 | 17 | 8 | 5 |
| A024 / L | 23 | 20 | 19 | 14 |
| A002 / L | 17 | 14 | 12 | 9 |
| A023 / R | 15 | 11 | 13 | 7 |
| A002 / R | 17 | 14 | 7 | 4 |
| A029 / L | 25 | 22 | 11 | 8 |
| A027 / L | 23 | 20 | 14 | 10 |
| A035 / R | 16 | 13 | 10 | 8 |
| A014 / L | 23 | 21 | 13 | 9 |

SNR80 / SNR60 = Signal to Noise Ratio for 80% / 60% correctly perceived words

The order of the experimental strategies was randomized before the first visit, and subjects were not informed which strategy they were listening to, to avoid any experimental bias. However, we do not rule out that subjects were able to differentiate between the strategies, since the experimental programs used different combinations of pulse patterns to stimulate the implant, with the MP program being most similar to their clinical program.

### 2.6 Statistics

The analyses were performed in MATLAB R2023b (Version 23.2.0) and R Studio (Version 2023.09. 1+494). One-way ANOVA and the Tukey test were used for pairwise-comparisons of the experimental programs and the raw sentence perception and vowel identification scores and focused benefit scores [37]. Pearson’s correlation coefficients and linear regression was used to assess the relation between speech results and subject demographics and focused threshold parameters. Linear mixed effects models were used to measure the interaction of demographics and focused threshold parameters across the experimental programs. Individual t-tests with Benjamini-Hochberg [38] p-values adjustments to decrease false discovery rates were used for the comparison between clinically CNC-based grouped outcomes.

## 3 Results

The speech perception thresholds and vowel identification performance outcomes of the three experimental programs are shown as percentage correct score (y-axis) in Fig. 3 The top panels show the outcomes for IEEE sentences in quiet and noise, the bottom panels show the vowel identification scores. The boxes represent the grouped outcomes of all participants with the group median scores, and inter-quartile ranges, overlayed by the individual outcomes. The panels are divided into three subsections (x-axis), representing the experimental programs (MP, Mixed, and DT).

**Figure 3:**
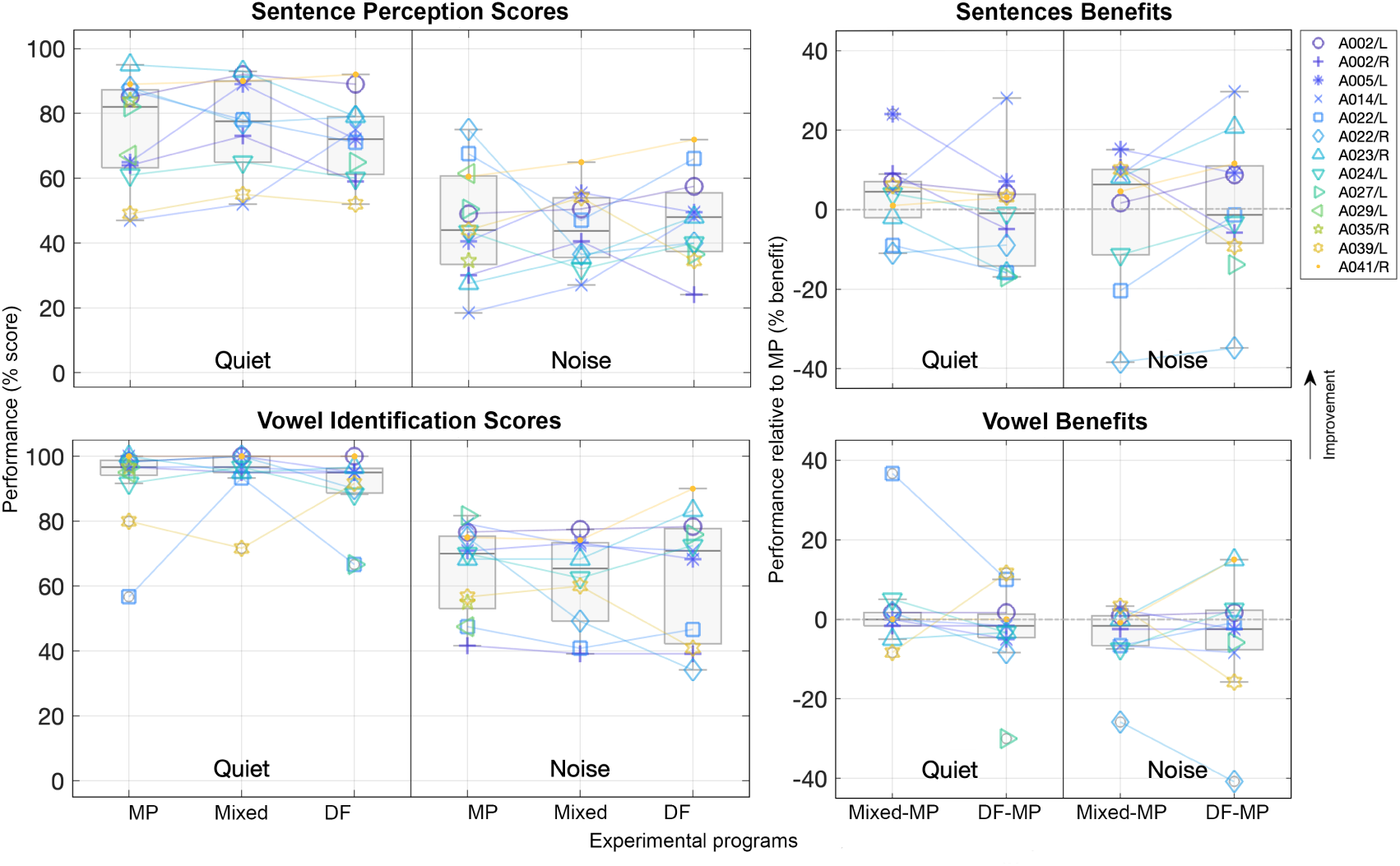
Speech perception (top) and vowel identification scores (bottom) for all participants. Data of all experimental programs, MP (n = 13), Mixed (n = 10), and DF (n = 11), is represented as boxes overlayed with the individual performance outcomes. The x-axis represents quiet and noisy listening conditions, grouped for the experimental programs, and the y-axis shows the performance scores in percent correct words; Performance benefits relative to MP (y-axis, in percent benefit) for Mixed program minus MP and DF minus MP (x-axis) for sentences (top right) and vowels (bottom right). Boxes represent the grouped performance benefits, and overlayed lines show the individual data.

Sentence performance scores in quiet (Fig. 3, top left), resulted in higher median percentage of correct words than sentences in noise, however comparable inter-quartile ranges (IQRs). In quiet, the MP program shows best outcomes (82% median, and IQR from 63.25% to 87.25% performance), followed by the Mixed program (77.5% median, IQR from 65% to 90%), and the DF program (72% median, IQR from 61.25% to 79%). Sentence performance scores in noise showed the highest median scores for the DF program (48% median, IQR from 37.38% to 55.5%), followed by the MP program (44% median, IQR from 33.38% to 60.75%), and slightly lower outcomes for the Mixed program (43.75% median, IQR from 35.5% to 54%).

Vowel identification scores (Fig. 3, bottom left) showed similar trends as seen for sentences, with decreasing performance scores from quiet to noise conditions. Generally, vowel identification in quiet shows ceiling effects in all three experimental programs. Median vowel outcomes in noise were highest for the DF program (70.83% median, and IQR of 42.29% and 77.71%), and MP (70% median, and IQR of 53.12% and 75.42%), and lowest with the Mixed program (65.42% median, and IQR of 49.17% and 73.33%).

Pairwise comparison showed that quiet performance across all programs was significantly different than performance in noise and for both sentences (larges group difference between the Mixed quiet and Mixed noise condition, p = .000104) and vowels (larges group difference between the same Mixed listening condition, p = .000015). Overall, no significant group differences were observed in performance in background noise, however, the DF program shows slightly better performance than the MP and Mixed programs for both sentences and vowels. Overall, individual across-strategy outcomes show large variability for sentence and vowels, except for the quiet listening situations, where several subjects were at ceiling (> 90%).

Performance benefits of the Mixed and DF programs over the MP are shown for sentences (Fig. 3, top right) and vowels (Fig. 3, bottom right). The benefit of the two programs over the MP program (Mixed-MP and DF-MP) are calculated by subtracting the MP percentage scores from the other scores. No benefit would be represented by zero percent on the y-axis.

Sentences in quiet show larger benefits for Mixed-MP (4.5% median, IQR between -2% and 7%) than for DF-MP (-1% median, IQR between -14.25% and 3.75%). Sentences in noise show similarly high median benefits for Mixed-MP (6.22% median, IQR between -11.5% and 10%) over DF-MP (-1.5% median, IQR between -8.63% and 10.79%). Grouped vowel benefits in quiet show less variations due to ceiling, with no significantly different benefit scores between Mixed-MP (0% median, IQR between -1.66% and 1.66%) and DF-MP (-1.67% median, IQR between -4.58% and 1.25%). Similar outcomes are observed for vowel benefits in noise with Mixed-MP (-1.67% median, IQR between -6.67% and 0.83%) and DF-MP (-2.5% median, IQR between -7.71% and 2.29%). Although there is no consistent trend that all participants do better with the DF or Mixed programs, seven out of ten individuals benefit from some degree of focusing for sentences in quiet and noise. One individual (A022) consistently performed lower in all focused programs than MP with both ears. A023/R did not show sentence benefits in quiet, and A024/L in noise. The largest average increase in benefit is observed for A014/L in quiet with an increase of 23 percentage points from 5% to 28% between Mixed-MP and DF-MP. Largest decrease is observed for A005/L with -17 percentage points from 24% to 7%. In noise, the majority of seven subjects show larger benefits of DF-MP over Mixed-MP (A014/L, A022/L, A022/R, A023/R, A024/L, A041/R) with maximal benefit scores of 24 for A014/L.

The ceiling effects for vowel in quiet limit the analyses of those data. Hence, most of the subjects show only small variable differences between 5% and -8.33% in Mixed-MP and between 2% and -8.33% in DF-MP, and overall slight decreases, similarly to the sentence benefits in quiet. Two subjects (A022/L, A039/L) deviate largely from those findings, with reversed trends, an effect that can be traced back to the raw performance data, presented in Fig. 3 (bottom left). Vowel benefits in noise show increasing trends from Mixed-MP to DF-MP for half of the subjects (A002/L, A022/L, A023/R, A024/L, A041/R) and maximal benefit of 15.8 percentage points for A041/R.

To better characterize the individual difference in raw performance and performance benefit with Mixed and DF programming compared to MP, we explored several demographic factors, duration of CI experience (Fig. 4), age (Fig. 5), and clinical performance based on CNC word scores (Fig. 6) and (Fig. 7). Vowel scores in quiet were excluded due to the observed ceiling effects. Fig. 4 shows the speech performance outcomes (left) and benefit calculations (right) in comparison to the duration of CI use (in years). The focused programs show larger deviations from the MP program for sentences and vowels in noise, such that performance is better with some focusing for participants with shorter duration of CI experience. Pearson’s correlation coefficients for sentence performance scores in noise with the MP program show a moderate correlation with CI experience (r = .59, *p* = .03), and slightly negatively correlated Mixed scores for vowels in noise (r = -.66, *p* = .04). DF outcomes show a strong trend, but do not reach statistical significance (r = -.59, *p* = .06). The observed increasing trend of MP programming for sentences in quiet in noise and CI experience compared to DF programming is somewhat expected, originating in the similarity of this programming approach to the subjects’ everyday clinical program. Benefit outcomes for the focused programs show decreasing trends with increasing CI use. The strongest relationships are found for Mixed-MP benefits, for both sentences (r = -.75, *p* = .01) and vowels (r = -.66, *p* = .04) in noise. DF-MP benefits show similar but not significant trends for sentences in noise (r = -.45, *p* = .17) and vowels in noise (r = -.43, *p* = .19), and also both focused programs showed the same trends for sentences in quiet, MF-MP (r = -.50, *p* = .14), and DF-MP (r = -.30, *p* = .37). Overall, shorter CI experiences consistently show larger focused program benefits over the MP program, especially for sentences in quiet and background noise.

**Figure 4:**
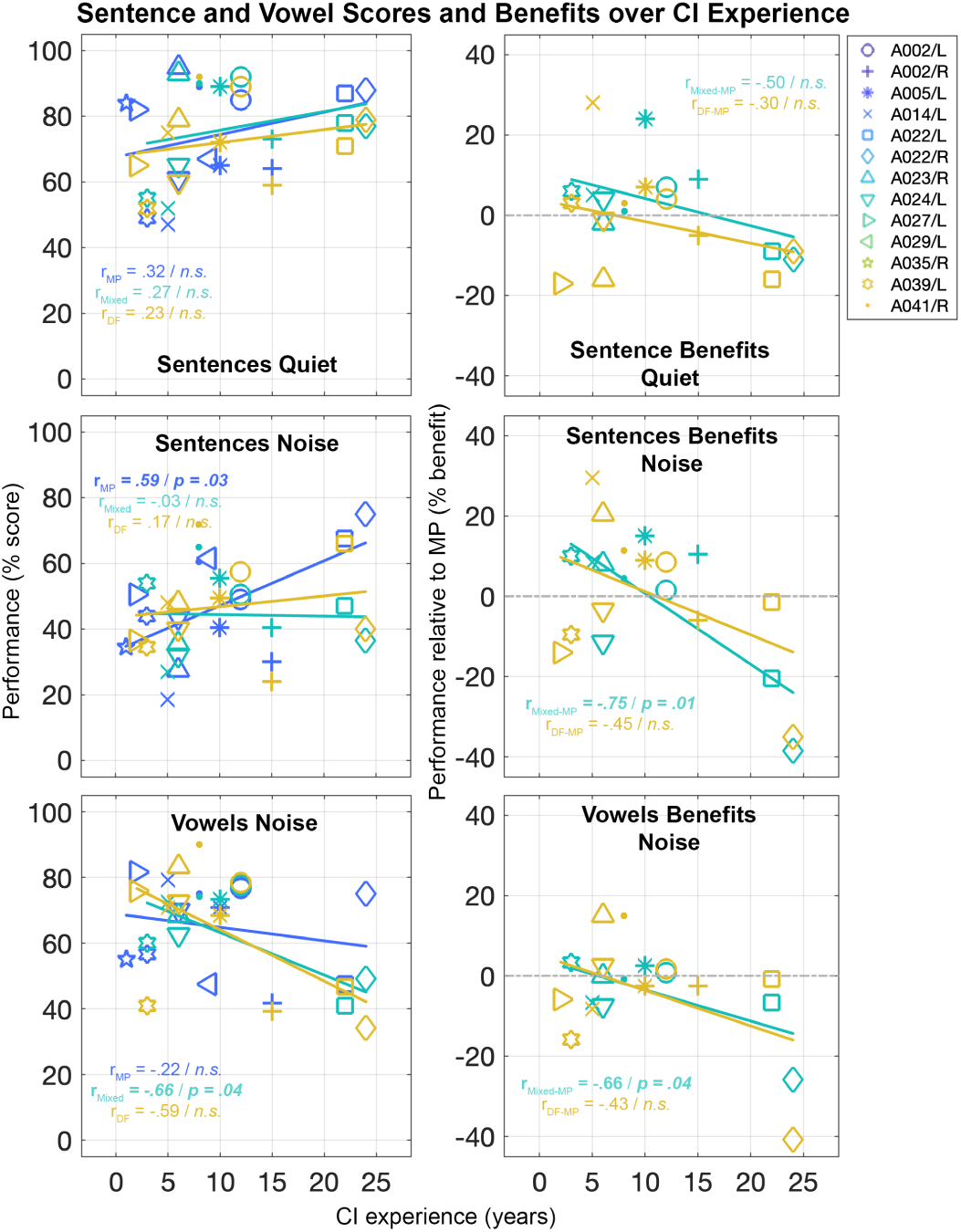
Sentence and vowel outcomes (y-axis, in percent correct words) over CI experience (x-axis, in years) for all experimental programs (color coded) and sentences in quiet and noise, and for vowels in noise (left), and Mixed (n = 10) and DF (n = 11) performance relative to MP (y-axis, in percent benefit) over CI experience for the same listening conditions (right).

**Figure 5:**
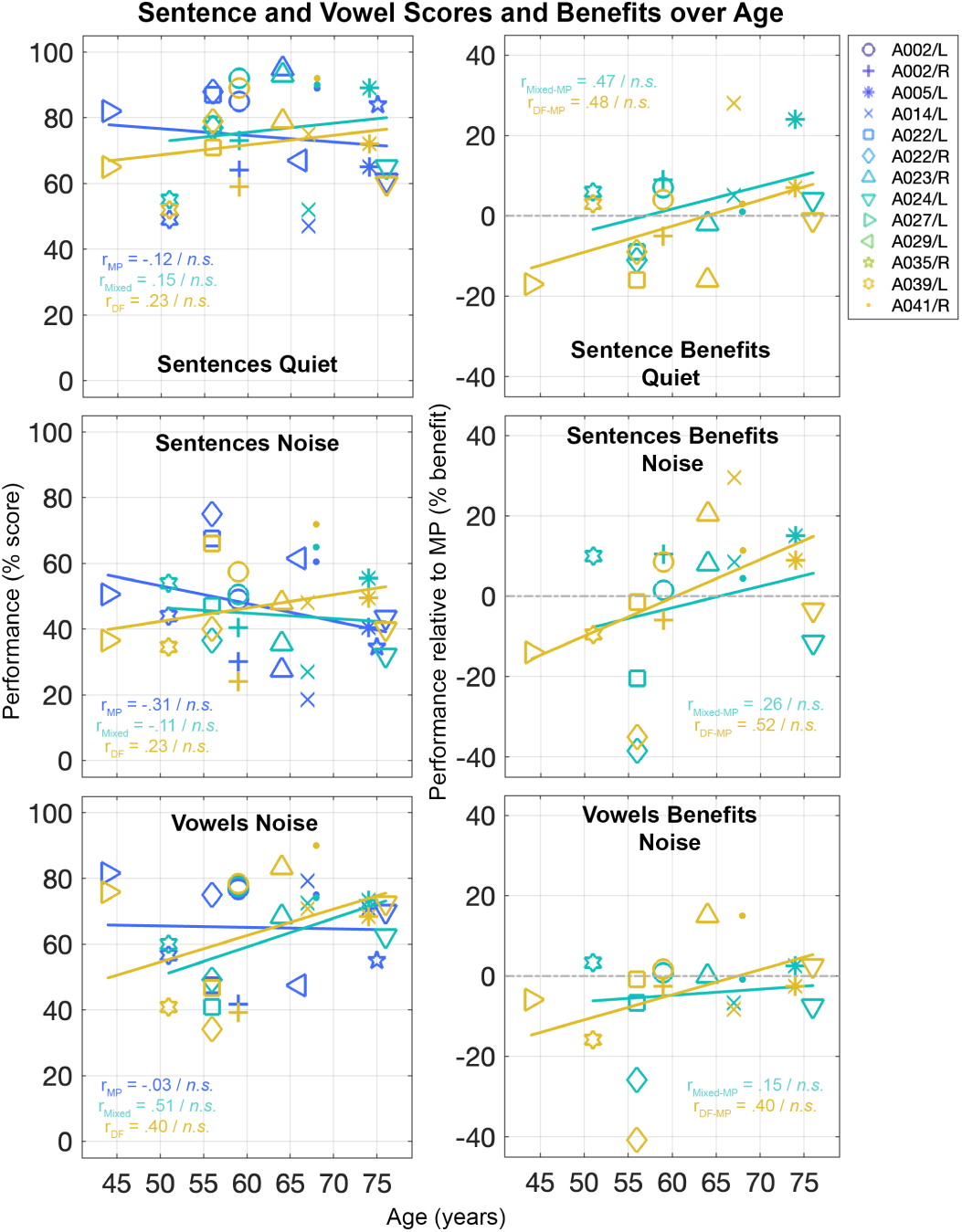
Sentence and vowel outcomes (y-axis, in percent correct words) over age (x-axis, in years) for all experimental programs (color coded) and sentences in quiet and noise (left), and Mixed (n = 10) and DF (n = 11) performance relative to MP (y-axis, in percent benefit) over age for the same listening conditions (right).

**Figure 6:**
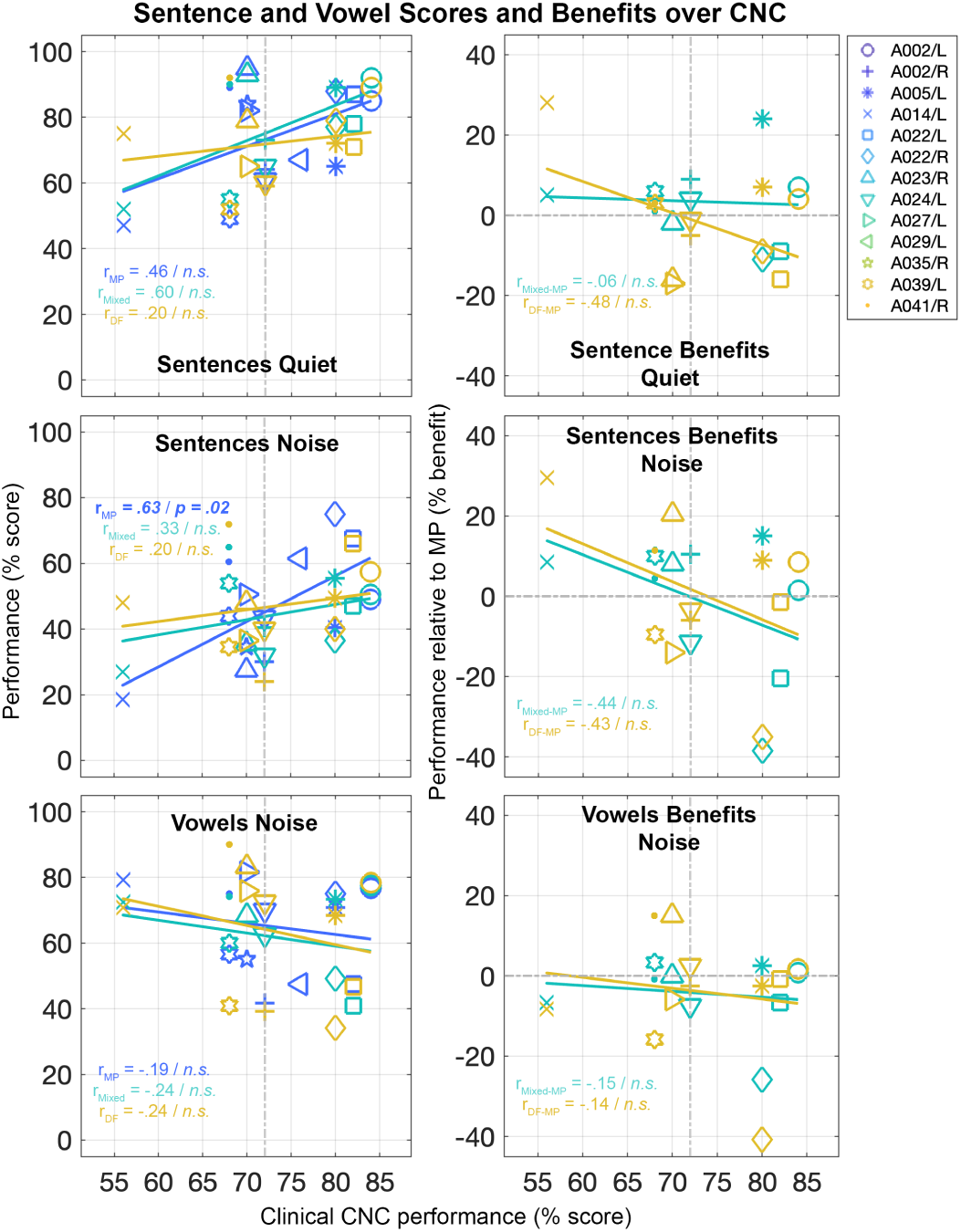
Sentence and vowel outcomes over clinical CNC performance (x-axis and y-axis, in percent correct words) for all experimental programs (color coded) and sentences in quiet and noise, and for vowels in noise (left), and Mixed (n = 10) and DF (n = 11) performance relative to MP (y-axis, in percent benefit) over clinical CNC performance for the same listening conditions (right). Median CNC performance marked as vertical line at 72% performance

**Figure 7:**
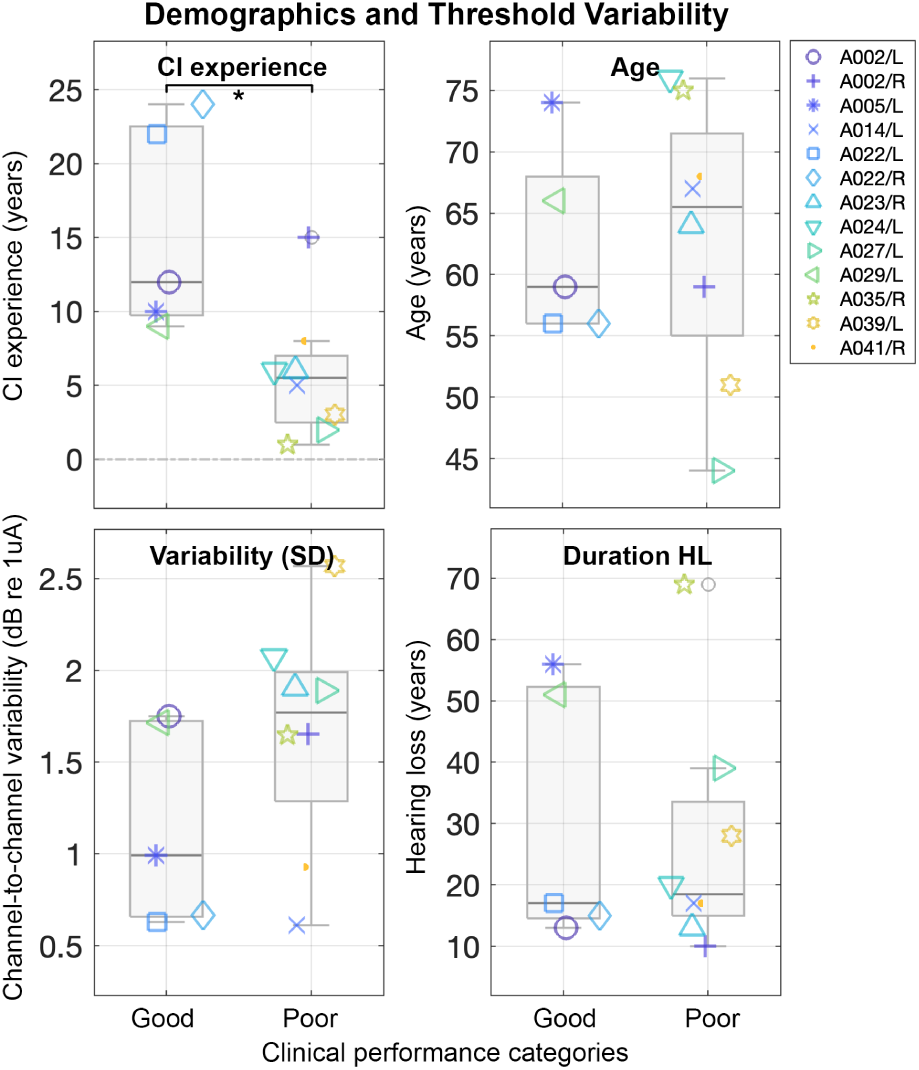
Median CNC split (at 72% performance) into good and poor performers and subject demographics (CI experience, age, duration of hearing loss) and focused detection threshold vari-abilities (in dB re 1 µA).

Fig. 5 shows the effect of age on speech performance (Fig. 5, left) and benefits (Fig. 5, right) scores. A possible co-linearity between age and CI experience was ruled out based on a low Pearson’s r and not significant *p*-value (r = -.21, *p* = .5). Speech performance outcomes (left) and benefit calculations (right) are presented in comparison to age (in years). Linear regression lines of speech perception scores as a function of age show slightly decreasing trends for the MP program for sentences in noise (r = -.31, *p* = .30), a trend we also observed in the clinical CNC scores and increasing age with the subjects’ everyday programs. On the contrary, increasing benefits of both focused programs and sentences in quiet, Mixed (r = .47, *p* =.17) and DF (r = .48, *p* = .14), and noise, Mixed (r = 26, *p* = .47) and DF (r = .52, *p* = .10), and the DF program for vowels in noise (r = .40, *p* = .23), suggesting that older participants were more likely to benefit from focused strategies, although the trends did not reach significance.

Bierer and Litvak [2] showed that clinically good and poor performers showed differences in benefits obtained with channel deactivation and current focusing. We, therefore, investigated this by correlating speech performance (Fig. 6, left) and benefits (Fig. 6, right) scores to the study subjects’ clinical CNC performance scores. Only the MP program showed a significant trend for sentences in noise over CNC scores (r = .63, *p* = .02). The Mixed program performance showed the strongest relationship as a function of clinical performance in the sentence in quiet condition (r = .60, *p* = .07) over the MP (r = .46, *p* = .11) and the DF program (r = .20, *p* = .56). All programs performed similarly in the vowel in noise listening condition. The DF program shows slight benefits for sentences in quiet (r = -.48, *p* = .13 ), and benefits are also observed for both focused programs for sentences in noise, MF (r = -.44, *p* = .21) and DF (r = -.43 , *p* = .18), with higher benefits for participants with lower CNC scores, indicating a similar, however, not significant trend as it was observed in previous work. Poor clinical performers tend to improve for focused programming, while good performers did not.

Similarly to Bierer and Litvak [2] and based on the observations in Fig. 6, we split subjects of the current study into two groups based on the median clinical recorded CNC performance of 72% (poor performers ď 72% performance, good performers ą 72% performance), marked in Fig. 6. Fig. 7 shows the separation of good and poor performers and subject demographics and focused threshold channel-to-channel variability. Individual t-tests and corrected p-values showed that CI duration significantly differs between good and poor clinical performers (*p* = .011), and no significance can be found for age, similar as shown before in the correlations. Previous studies identified slight differences of duration of hearing loss and for channel-to-channel variations between good and poor performers, trends that were also observed in the current study cohort. We also investigated differences between good and poor performers in sentence and vowel benefit scores of the Mixed and the DF programs relative to MP (Fig. 8), however, there were no additional significant differences aside from what we found for CI duration.

**Figure 8:**
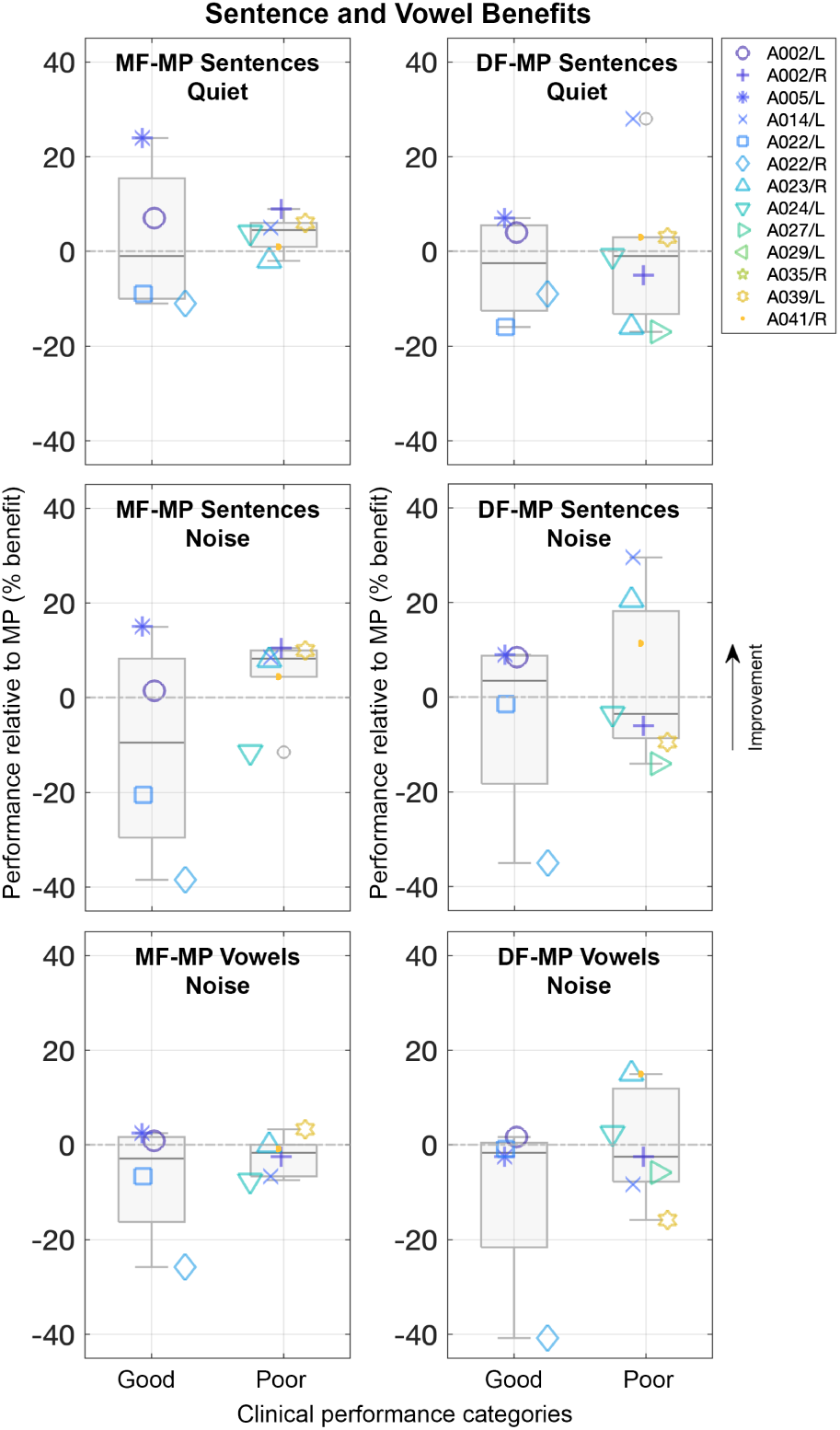
Median CNC split (at 72% performance) into good and poor performers and sentence and vowel performance benefits for Mixed and DF programs over the MP program.

Due to the large variabilities observed across listening conditions in the MP, Mixed, and DF programs, linear mixed effects models were used to test for interaction effects of CI duration, age, clinical CNC scores and variability of focused thresholds versus experimental programs on sentence in noise scores. Subject ID was included as a random intercept to account for repeated measures from each subject. The MP program was coded as the reference condition in each model. A significant interaction between programming conditions was only present for CI experience (*p* = .036). Interaction effects of age, and CNC score, and focused thresholds channel-to-channel variability did not reach significance. Much of the variance across conditions and subjects was unaccounted for, with marginal and conditional *R*^2^ of the CI experience model reaching only 0.167 and 0.518, respectively.

Our previous work controlled rates and pulse widths across subjects and focused programs [2, 33]. Therefore, to rule out the potential influence of the automatically optimized stimulation rate and pulse widths of MP, Mixed, and DF programs, we investigated the programming differences as a function of speech perception outcomes, similarly as shown for the correlations of demographics and the raw performance scores. Due to the increased power consumption of focused stimulation, the automatically adjusted pulse widths tend to increase for focused programs compared to the MP program, and the rates decrease (Table 2). Correlations of speech performance as a function of pulse width and rate did not show significant differences in any of the experimental programs throughout listening conditions. However, we found a similar but not significant difference between good and poor performers with smaller pulse widths and rate differences between focused programs and the MP programming.

## 4 Discussion

The overall rationale of this study and of optimizing individual programs is the reduction of unintended neural activation or channel interaction caused by electric stimulation and, therefore, to improve speech perception outcomes. By combining two previously explored concepts, channel deactivation and current focusing, the current work investigated both factors through a series of different programs to quantify the impacts on sentence and vowel scores, and the influence of related demographics [2, 33].

With the ultimate goal of reducing channel interactions and improving speech perception outcomes, we automatically selected subsets of channels for deactivation and combined this with current focusing approaches. To investigate the effects of different stimulation settings on the remaining active electrodes, the current algorithm was further optimized to identify candidates of MP and DT channels based on focused perception thresholds.

By calculating the relative performance of DF and Mixed programs to the MP program (Fig. 3, top right), our outcomes show only small trends for the focused programs over the MP program for sentences in quiet and noisy listening conditions on a group level, however, most subjects benefit from some degree of focusing for sentences in quiet and in noise. Mixed programming shows larger benefits over the MP than DF over MP for most of the listeners for sentences in quiet, and most individuals seem to benefit even more from DF over MP than for Mixed over MP programming and sentences in noise. This might be due to the fact, that the Mixed program comprises a combination of MP and DF channels, which makes the overall programming more akin to the individuals’ everyday strategy. While this effect benefits in a quiet listening condition, the partially defined MP channels might create larger disadvantageous channel interactions in noise, covering the benefits of the individually focused channels. Overall, increased focusing counteracts broad unintended neural activation, hence, larger listening benefits are achieved.

The improvement with CI experience observed with the MP program (Fig. 4, left) is an expected trend due to the similarity of the MP program to participants’ clinical coding strategy. Conversely, focused programs resulted in higher benefits for participants with CI experience less than about 5 to 10 years (Fig. 4, right). This may be a result of long durations of acclimatization to their MP clinical strategy limiting the ability of adaptation to new unfamiliar CI programming. Duration of deafness, respectively, duration before being implanted, was found to negatively correlate (r = -.36, *p* = 0.16) with performance scores in speech perception [33] before. It can be seen in Table 1, that the current cohort of study participants indeed includes some individuals with long durations of hearing loss. Nevertheless, the current outcomes could not reproduce this previous relationship, however, showed a positive correlation for benefits of DF over MP programming for sentences and vowels in noise with older age instead (Fig. 5, right). The linear regressions as function of age indicated slight improvements for sentences in noise around approximately 60 years of age. Similarly, vowel in noise benefits of DF over MP showed a positive trend, however, all trends did not reach statistical significance. The current age effects might be relatable to the previous trends for the duration of deafness, to some extent, via shorter durations of CI experience in older individuals, and vice versa, supported by the observed trends of these two demographics. In conclusion, optimal CI programming may need to occur before participants have decades of CI experience to achieve maximal individual performance abilities with these strategies.

The demographic differences of duration of CI experience, age, and duration of hearing loss between good and poor performers are shown in Fig. 7. The split is based on the clinical CNC performance, which indicated a decreasing trend in benefits of poor performing individuals, respectively, lower CNC scores (Fig. 6, bottom). In contrast to previous work, duration of deafness is similarly distributed between good and poor performers. However, the duration of CI experience and age effects are again visible and in line with our investigations across all study subjects, showing that good performers are generally older, however, with large variabilities across subjects. One possible explanation of the across-subject performance differences might be the underlying detection thresholds. Individually focused threshold profiles are highly variable, hence, the optimized channel selection varied across listeners, similar to previous findings [2, 25, 31]. A reduction in the overall focused threshold channel-to-channel variability might be beneficial for those subjects who have large variability, a possible effect of variations in neural survival, and the quality of the ENI. In the current study, high-focused threshold channels were deactivated to reduce the variability, and the remaining channels were set to MP, DF, or Mixed stimulation. We only found a small and not significant effect of higher channel-to-channel variabilities in poor compared to good performers (Fig. 7), however, a higher number of deactivated channels for those individuals might be a possible indicator for successfully improving channel frequency selectivity and the corresponding performance, especially by deactivating high focused threshold channels to optimize the transmission accuracy at the ENI.

As presented in the current study, sentence and vowel identification scores in background noise seem to increase for the experimental DF program with deactivated channels compared to the tested MP program. Bierer and Litvak [2] and Arenberg et al. [33] used an algorithm to select candidates for channel deactivation based on focused threshold profiles, a predecessor of the algorithm presented in this work. Deactivating channels with large channel interactions might reduce unwanted neural activations and consequently improve both vowel identification and sentence recognition abilities. The current findings might show the potential improvements because of reduced channel interactions, and improved spectral resolution through a more precise signal transmission at the ENI. Vowels carry crucial features for speech perception, such as first and second formant (*F1*, *F2* ) and related higher-order frequency information, and are, in addition, reliant to spectral cues, even in noisy listening situations [39, 40]. Increasing the formant accuracy and vowel formant contrasts, respectively, increasing spectral content via channel deactivation and current focusing, might benefit the speech perception outcomes in background noise [41, 42]. Even though the variability of the subject group is high, and no significance can be reached for most of the observed trends, the results of this study show the same trend compared to previous work [28, 29, 33]. The slightly different trends might be accountable to the differences in the used materials. The investigation of spondees did not reveal any benefits [33], the Hearing in noise test (HINT) and Monosyllables (Berenstein 2008, Srinivasan 2013), on the contrary, showed benefits of focused stimulation. The overall similarity in benefits between Mixed and DF over MP programs for sentences and vowels in background noise with the low context IEEE speech material used in this study might have been a good match to the complexity of the vowel identification task, where the CI listeners cannot rely on the presented context to score the correct words.

Clinically, electrode deactivation is a commonly used and often-necessary method to avoid unwanted electric stimulation effects, such as facial stimulation. In certain cases, the location of the implanted electrode array penetrates cochlea tissue or is affected by electrode tip fold-overs, leading to suboptimal stimulation locations and effects, e.g. [10]. In other cases, stimulation intensities, respectively, current, on certain electrodes run compliance issues, which limits the presentation of sufficiently loud signals. In both cases, electrodes can be deactivated in the individual’s clinical programming. Aside from clinically necessary channel deactivation, there are other negatively influencing factors, that reduce performance outcomes. High thresholds and broad channel interaction, for example, were identified in previous studies as possible factors of decreased performance scores [6, 31, 43]. Such channels activate broad neural regions at the ENI, respectively, generate large channel interactions, which might negatively influence low threshold neighboring channels, hence, superimpose the beneficial narrow neural activation of those channels. Clinically, one way of avoiding interacting electric signals is achieved by channel activation patterns with larger distances between active channels. A recent study tried to counteract broad interactions by deactivating intermediate channels and alternating the selection to compensate for missing CI channels, and the outcomes of the study showed small benefits for sentences in noise [16]. Nevertheless, the broad activation resulting from MP stimulation might still negatively impact the accuracy of the signals along the electrode array. Identifying, deactivating, and focusing CI channels, that impact the listening benefit through stimulation inaccuracies as the result of large channel interactions or high focused thresholds of poor ENI channels, might in addition improve speech perception. Incorporating channel interaction measures in coding approaches were explored before, however, there is still a gap for clinical applications due to high individual variabilities across CI listeners.

Beside channel deactivation, the current study also explored activating the remaining CI channels with a mix of DT and MP stimulation. Overall, the Mixed configuration showed lower benefits, compared to the DF programming with deactivated channels, especially for sentences and vowels in background noise. Even though that there were no additional listening benefits of the Mixed channel configuration, one positive aspect of the combination of MP and DT channels in one program might be a potentially increased battery lifetime. Arenberg et al. [33] has already argued that the battery lifetime can be improved in TP stimulation by dynamically adjusting the amount of focusing and partly returning the currents to the external return electrode, resulting in less power required, respectively, reduced stimulation amplitudes to generate sufficiently loud signals. Hence, programming a subset of channels in MP mode might further reduce the charge needed. Unfortunately, we could not assess the battery lifetime during the acute testing performed in this study, because the sessions were limited to maximal three hours per visit. Further, one previous study demonstrated the importance of channel selection for focused stimulation, such that performance significantly declined if channels were focused when they were close to the inner wall of the cochlea [44]. There is a chance that in this study, without the aid of CT imaging to assess the electrode location relative to the inner wall, that we may have selected suboptimally which channels to focus.

One limiting factor of the current study is the acute nature of the experiments. Bierer and Litvak [2] suggested exploring the long-term effects of channel deactivation. A later study by Arenberg et al. [33] also suggested monitoring performance over a longer duration, to acclimatize to the different stimulation patterns of the optimized programming. CI listeners have shown that over longer durations of adaptation (days up to several weeks) to frequency allocations on active channels can be compensated and speech perception improved [45, 46]. We may be underestimating the performance outcomes of CI participants, as they might adapt to deactivated channels and DT stimulation if provided sufficient experience. This is supported by the evidence that we observed with DT and Mixed strategies related to the duration of CI experience, suggesting the importance of acclimatization to new programming strategies. Therefore, we are currently investigating the long-term effects of our programming approaches, with a take-home trial to determine the overall effects our strategy after listening in the real world.

In summary, despite the large variability in performance outcomes with the three experimental strategies, CI individuals showed some improvements in speech perception and vowel identification tasks in quiet and background noise utilizing optimized Mixed and DT programming incorporating deactivated high threshold channels, compared to MP programming with the same deactivated channels. In addition, poor performers, older listeners, and listeners with shorter CI experience seem to benefit more from program optimization that included focused stimulation (Mixed and/or DT). Furthermore, clinically poor performers are more likely to have higher focused detection threshold variability, a possible indicator for poor quality of the underlying ENI. Future studies will work to further optimize the automatic algorithm for selecting channels for deactivation and DT stimulation. We are currently working on the validation of an extended automatic channel selection algorithm that takes large variability in focused detection threshold profiles as well as the degree of asymmetry into account (e.g. threshold tilt and average values), to further optimize channel selection.

## Acknowledgements

The authors would like to thank the cochlear implant participants for their time and effort in the study and their patience in performing the experiments. The authors also want to acknowledge Scott Aker for supporting the data collection, Faten Awwad for the participant recruitment, and Yan Zhao for assistance with statistical analyses. This work was funded by the National Institutes of Health, National Institute of Deafness and Other Communication Disorders R01DC012142 (J.G.A) The authors declare no conflicts of interest.

## References

[1] Fan-Gang Zeng. Celebrating the one millionth cochlear implant. JASA Express Letters, 2(7): 077201, 2022. doi: 10.1121/10.0012825.

[2] Julie A. Bierer and Leonid Litvak. Reducing Channel Interaction Through Cochlear Implant Programming May Improve Speech Perception. Trends in Hearing, 20(20):1–12, 2016. ISSN 23312165. doi: 10.1177/2331216516653389.

[3] Belinda A. Henry and Christopher W. Turner. The resolution of complex spectral patterns by cochlear implant and normal-hearing listeners. The Journal of the Acoustical Society of America, 113(5):2861–2873, 2003. ISSN 0001-4966. doi: 10.1121/1.1561900.

[4] Belinda A. Henry, Christopher W. Turner, and Amy Behrens. Spectral peak resolution and speech recognition in quiet: Normal hearing, hearing impaired, and cochlear implant listeners. The Journal of the Acoustical Society of America, 118(2):1111–1121, 2005. ISSN 0001-4966. doi: 10.1121/1.1944567.

[5] Jong Ho Won, Ward R. Drennan, and Jay T. Rubinstein. Spectral-ripple resolution correlates with speech reception in noise in cochlear implant users. JARO -Journal of the Association for Research in Otolaryngology, 8(3):384–392, 2007. ISSN 15253961. doi: 10.1007/s10162-007-0085-8.

[6] Gary L. Jones, Jong Ho Won, Ward R. Drennan, and Jay T. Rubinstein. Relationship between channel interaction and spectral-ripple discrimination in cochlear implant users. The Journal of the Acoustical Society of America, 133(1):425–433, 2013. ISSN 0001-4966. doi: 10.1121/1. 4768881.

[7] N. R.A. Van Groesen, J. J. Briaire, and J. H.M. Frijns. Evaluation of Two Spectro-Temporal Ripple Tests and Their Relation to the Matrix Speech-in-Noise Sentence Test in Cochlear Implant Recipients. Ear and Hearing, 44(5):1221–1228, 2023. ISSN 15384667. doi: 10.1097/AUD.0000000000001365.

[8] Jack H. Noble, Reńe H. Gifford, Andrea J. Hedley-Williams, Benoit M. Dawant, and Robert F. Labadie. Clinical evaluation of an image-guided cochlear implant programming strategy. Audiology and Neurotology, 19(6):400–411, 2014. ISSN 14219700. doi: 10.1159/000365273.

[9] Ning Zhou. Deactivating stimulation sites based on low-rate thresholds improves spectral ripple and speech reception thresholds in cochlear implant users. The Journal of the Acoustical Society of America, 141(3):EL243–EL248, 2017. ISSN 0001-4966. doi: 10.1121/1.4977235.

[10] Fabiana Danieli, Thomas Dermacy, Maria Stella Arantes do Amaral, Ana Cĺaudia Miran-dola Barbosa Reis, Dan Gnansia, and Miguel Angelo Hyppolito. Auditory performance of post-lingually deafened adult cochlear implant recipients using electrode deactivation based on postoperative cone beam CT images. European Archives of Oto-Rhino-Laryngology, 278 (4):977–986, 2021. ISSN 14344726. doi: 10.1007/s00405-020-06156-8.

[11] Sarah E. Warren and Samuel R. Atcherson. Evaluation of a clinical method for selective electrode deactivation in cochlear implant programming. Frontiers in Human Neuroscience, 17, 2023. ISSN 16625161. doi: 10.3389/fnhum.2023.1157673.

[12] Deborah Vickers, Aneeka Degun, Angela Canas, Thomas Stainsby, and Filiep Vanpoucke. Deactivating cochlear implant electrodes based on pitch information for users of the ACE strategy. Advances in Experimental Medicine and Biology, 894:115–123, 2016. ISSN 22148019. doi: 10.1007/978-3-319-25474-6 13.

[13] Joke A. Debruyne, Tom Francart, A. Miranda L. Janssen, Kim Douma, and Jan P.L. Brokx. Fitting prelingually deafened adult cochlear implant users based on electrode discrimination performance. International Journal of Audiology, 56(3):174–185, 2017. ISSN 17088186. doi: 10.1080/14992027.2016.1243262.

[14] G. E. Peterson and I. Lehiste. Revised CNC lists for auditory tests. The Journal of speech and hearing disorders, 27:62–70, 1962. ISSN 00224677. doi: 10.1044/jshd.2701.62.

[15] Anthony J. Spahr, Michael F. Dorman, Leonid M. Litvak, Susan Van Wie, Rene H. Gifford, Philipos C. Loizou, Louise M. Loiselle, Tyler Oakes, and Sarah Cook. Development and validation of the azbio sentence lists. Ear and Hearing, 33(1):112–117, 2012. ISSN 01960202. doi: 10.1097/AUD.0b013e31822c2549.

[16] D. M. Wohlbauer, Wai Kong Lai, and Norbert Dillier. InterlACE Sound Coding for Unilateral and Bilateral Cochlear Implants. IEEE Transactions on Biomedical Engineering, 71 (3):904–915, mar 2023. ISSN 0018-9294. doi: 10.1109/TBME.2023.3322348. URL https://ieeexplore.ieee.org/document/10272661/.

[17] Andrej Kral, Rainer Hartmann, Dariusch Mortazavi, and Rainer Klinke. Spatial resolution of cochlear implants: The electrical field and excitation of auditory afferents. Hearing Research, 121(1-2):11–28, 1998. ISSN 03785955. doi: 10.1016/S0378-5955(98)00061-6.

[18] Julie Arenberg Bierer and John C. Middlebrooks. Auditory cortical images of cochlear-implant stimuli: Dependence on electrode configuration. Journal of Neurophysiology, 87(1):478–492, 2002. ISSN 00223077. doi: 10.1152/jn.00212.2001.

[19] Julie Arenberg Bierer and John C. Middlebrooks. Cortical Responses to Cochlear Implant Stimulation: Channel Interactions. JARO -Journal of the Association for Research in Otolaryngology, 5(1):32–48, 2004. ISSN 15253961. doi: 10.1007/s10162-003-3057-7.

[20] Russell L. Snyder, Julie A. Bierer, and John C. Middlebrooks. Topographic spread of inferior colliculus activation in response to acoustic and intracochlear electric stimulation. JARO -Journal of the Association for Research in Otolaryngology, 5(3):305–322, 2004. ISSN 15253961. doi: 10.1007/s10162-004-4026-5.

[21] Ben H. Bonham and Leonid M. Litvak. Current focusing and steering: Modeling, physiology, and psychophysics. Hearing Research, 242(1-2):141–153, 2008. ISSN 03785955. doi: 10.1016/ j.heares.2008.03.006.

[22] Julie G. Arenberg, Shigeto Furukawa, and John C. Middlebrooks. Auditory cortical images of tones and noise bands. JARO -Journal of the Association for Research in Otolaryngology, 1 (2):183–194, 2000. ISSN 15253961. doi: 10.1007/s101620010036.

[23] Monita Chatterjee. Effects of stimulation mode on threshold and loudness growth in multielectrode cochlear implants. The Journal of the Acoustical Society of America, 105(2):850–860, 1999. ISSN 0001-4966. doi: 10.1121/1.426274.

[24] Julie Arenberg Bierer and Amberly D. Nye. Comparisons between detection threshold and loudness perception for individual cochlear implant channels. Ear and Hearing, 35(6):641–651, 2014. ISSN 15384667. doi: 10.1097/AUD.0000000000000058.

[25] Julie Arenberg Bierer. Threshold and channel interaction in cochlear implant users: Evaluation of the tripolar electrode configuration. The Journal of the Acoustical Society of America, 121 (3):1642–1653, 2007. ISSN 0001-4966. doi: 10.1121/1.2436712.

[26] Lucas H.M. Mens and Carlo K. Berenstein. Speech perception with mono-and quadrupolar electrode configurations: A crossover study. Otology and Neurotology, 26(5):957–964, 2005. ISSN 15317129. doi: 10.1097/01.mao.0000185060.74339.9d.

[27] Chris van den Honert and David C. Kelsall. Focused intracochlear electric stimulation with phased array channels. The Journal of the Acoustical Society of America, 121(6):3703, 2007. ISSN 00014966. doi: 10.1121/1.2722047.

[28] Carlo K. Berenstein, Lucas H.M. Mens, Jef J.S. Mulder, and Filiep J. Vanpoucke. Current steering and current focusing in cochlear implants: Comparison of monopolar, tripolar, and virtual channel electrode configurations. Ear and Hearing, 29(2):250–260, 2008. ISSN 01960202. doi: 10.1097/AUD.0b013e3181645336.

[29] Arthi G. Srinivasan, Monica Padilla, Robert V. Shannon, and David M. Landsberger. Improving speech perception in noise with current focusing in cochlear implant users. Hearing Research, 299:29–36, may 2013. ISSN 03785955. doi: 10.1016/j.heares.2013.02.004.

[30] Ching Chih Wu and Xin Luo. Current steering with partial tripolar stimulation mode in cochlear implants. JARO -Journal of the Association for Research in Otolaryngology, 14(2): 213–231, apr 2013. ISSN 15253961. doi: 10.1007/s10162-012-0366-8.

[31] Christopher J. Long, Timothy A. Holden, Gary H. McClelland, Wendy S. Parkinson, Clough Shelton, David C. Kelsall, and Zachary M. Smith. Examining the electro-neural interface of cochlear implant users using psychophysics, CT scans, and speech understanding. JARO -Journal of the Association for Research in Otolaryngology, 15(2):293–304, 2014. ISSN 14387573. doi: 10.1007/s10162-013-0437-5.

[32] Leonid M. Litvak, Anthony J. Spahr, and Gulam Emadi. Loudness growth observed under partially tripolar stimulation: Model and data from cochlear implant listeners. The Journal of the Acoustical Society of America, 122(2):967–981, 2007. ISSN 0001-4966. doi: 10.1121/1.2749414.

[33] Julie G. Arenberg, Wendy S. Parkinson, Leonid Litvak, Chen Chen, Heather A. Kreft, and Andrew J. Oxenham. A dynamically focusing cochlear implant strategy can improve vowel identification in noise. Ear and Hearing, 39(6):1136–1145, 2018. ISSN 15384667. doi: 10.1097/AUD.0000000000000566.

[34] Benjamin Caswell-Midwinter and Julie G. Arenberg. Comparing Fixed and Individualized Channel Interaction Coefficients for Speech Perception With Dynamic Focusing Cochlear Implant Strategies. Trends in Hearing, 27, jan 2023. ISSN 23312165. doi: 10.1177/23312165231176157. URL https://journals.sagepub.com/doi/full/10.1177/ 23312165231176157.

[35] Johan H.M. Frijns, David M.T. Dekker, and Jeroen J. Briaire. Neural excitation patterns induced by phased-array stimulation in the implanted human cochlea. Acta Oto-Laryngologica, 131(4):362–370, 2011. ISSN 00016489. doi: 10.3109/00016489.2010.541939.

[36] Julie Bierer. Probing the Electrode-Neuron Interface With Focused Cochlear Implant Stimulation. Trends in Amplification, 14(2):84–95, 2010. ISSN 10847138. doi: 10.1177/1084713810375249.

[37] Olive Jean Dunn. Multiple Comparisons among Means. Journal of the American Statistical Association, 56(293):52–64, 1961. ISSN 1537274X. doi: 10.1080/01621459.1961.10482090.

[38] Benjamini Hochberg. Benjamini Hochberg1995.Pdf, 1995.

[39] James Hillenbrand, Laura A. Getty, Michael J. Clark, and Kimberlee Wheeler. Acoustic characteristics of American English vowels. The Journal of the Acoustical Society of America, 97 (5):3099–3111, may 1995. ISSN 0001-4966. doi: 10.1121/1.411872. URL /asa/jasa/article/ 97/5/3099/847219/Acoustic-characteristics-of-American-English.

[40] Rikus Swanepoel, Dirk J. J. Oosthuizen, and Johan J. Hanekom. The relative importance of spectral cues for vowel recognition in severe noise. The Journal of the Acoustical Society of America, 132(4):2652–2662, 2012. ISSN 0001-4966. doi: 10.1121/1.4751543.

[41] Philipos C. Loizou and Oguz Poroy. Minimum spectral contrast needed for vowel identification by normal hearing and cochlear implant listeners. The Journal of the Acoustical Society of America, 110(3):1619–1627, 2001. ISSN 0001-4966. doi: 10.1121/1.1388004.

[42] Taffeta M. Elliott and Fédéric E. Theunissen. The modulation transfer function for speech intelligibility. PLoS Computational Biology, 5(3), 2009. ISSN 15537358. doi: 10.1371/journal.pcbi.1000302.

[43] Elizabeth S. Anderson, David A. Nelson, Heather Kreft, Peggy B. Nelson, and Andrew J. Oxenham. Comparing spatial tuning curves, spectral ripple resolution, and speech perception in cochlear implant users. The Journal of the Acoustical Society of America, 130(1):364–375, 2011. ISSN 0001-4966. doi: 10.1121/1.3589255.

[44] Lindsay DeVries and Julie G. Arenberg. Psychophysical Tuning Curves as a Correlate of Electrode Position in Cochlear Implant Listeners. JARO -Journal of the Association for Research in Otolaryngology, 19(5):571–587, 2018. ISSN 14387573. doi: 10.1007/s10162-018-0678-4.

[45] Qian-Jie Fu, Robert V. Shannon, and John J. Galvin. Perceptual learning following changes in the frequency-to-electrode assignment with the Nucleus-22 cochlear implant. The Journal of the Acoustical Society of America, 112(4):1664–1674, 2002. ISSN 0001-4966. doi: 10.1121/1.1502901.

[46] John J. Galvin, Qian Jie Fu, and Geraldine Nogaki. Melodic contour identification by cochlear implant listeners. Ear and Hearing, 28(3):302–319, 2007. ISSN 01960202. doi: 10.1097/01.aud.0000261689.35445.20.

